# Epigenetic Effects of Exposure to Insecticide on Early Differentiation of Mouse Embryonic Stem Cells

**DOI:** 10.1101/628487

**Authors:** Wenlong Wang, Satoshi Otsuka, Hiroko Nansai, Tomohiro Ito, Kuniya Abe, Yoichi Nakao, Jun Ohgane, Minoru Yoneda, Hideko Sone

**Author notes:** Address correspondence to Hideko SONE, Center for Environmental Risk Research, National Institute for Environmental Studies, 16-2 Onogawa, Tsukuba, Ibaraki, 305-8606, Japan.

## Abstract

Increasing evidence indicates that insecticides induce various diseases via DNA methylation. DNA methylation plays an important role during cell differentiation and exhibits its greatest vulnerability to environmental factors during embryogenesis. Therefore, it is important to evaluate the effects on DNA methylation at the early stage of cell differentiation to understand developmental toxicity. However, DNA methylation induced by insecticides and the associated effects on cell differentiation are unclear. In this research, we introduced a high-content approach utilizing mouse embryonic stem cells harboring enhanced green fluorescent protein fused with methyl CpG-binding protein to evaluate global DNA methylation induced by various insecticides. DNA methylation was assessed in 22 genes after pesticide exposure to investigate the relationships with biological processes such as cell cycle, cell apoptosis, and cell differentiation. Exposure to acetamiprid, imidacloprid, carbaryl, and *o,p*′-DDT increased the granular intensity, indicating their global DNA-methylating effects. Exposure to imidacloprid decreased DNA methylation in genes such as Cdkn2a, Dapk1, Cdh1, Mlh1, Timp3, and Rarb, indicating the potential influence of the DNA methylation pattern on cell differentiation. We developed a promising approach for evaluating global DNA methylation, and our findings suggested that imidacloprid might exhibit developmental effects through DNA methylation pattern.

## Introduction

Insecticides exist in the environment, and they are associated with various adverse health outcomes such as neurological and reproductive disorders and cancer (Fritschi et al. 2015, Sanborn et al. 2007, Tiemann 2008, Zhang et al. 2016). Increasing evidence indicates that insecticides induce epigenetic effects as the toxicological mechanism (Costa 2015, Hodges et al. 2000). Recently, the evaluation of developmental toxicity has proven important (Hass 2006), and it was reported that epigenetic abnormalities are related to mammalian development such as brain development, particularly during early embryonic and germ cell development (Doherty and Roth 2016, Ideta-Otsuka et al. 2017, Keverne et al. 2015, Marczylo et al. 2016), indicating a need to evaluate the epigenetic effects of insecticides and their roles in developmental toxicity.

DNA methylation, as an important mechanism of epigenetic effects, regulates gene expression to control differentiation in many self-renewing tissues such as germline and hematopoietic stem cells (Klose and Bird 2006). For instance, global DNA methylation increases with the loss of methylation of specific genes that define cell identity during stem cell differentiation (Suelves et al. 2016). Decreased global and gene-specific DNA methylation commonly occurs in carcinogenesis (Teschendorff et al. 2014). Therefore, the evaluation of both global and gene-specific DNA methylation provides important insights into developmental toxicity and the associated toxicities of environmental chemicals. Evidences indicated that insecticides induce developmental neurotoxicity (Declerck et al. 2017) and carcinogenesis (Zhang et al. 2012) through changes in DNA methylation patterns. DNA methylation exhibits its greatest vulnerability to environmental factors during embryogenesis, (Foley et al. 2009, Relton and Davey Smith 2010), resulting in adverse health outcomes in children. Thus, DNA methylation during cell differentiation can provide an important insight to understand the developmental toxicity of insecticides.

Embryonic stem cells (ESCs) represent a powerful biological model for evaluating developmental toxicities (Tandon and Jyoti 2012). Moreover, their differentiation into normal somatic cells such neuronal (Okabe et al. 1996) and heart cells (Chen et al. 2010) permits assessments of diverse developmental toxicities such as developmental neurotoxicity. Differentiation-related genes have been used as endpoints for development toxicity tests (van Dartel et al. 2010), and the differentiation of ESCs can be controlled via DNA methylation (Altun et al. 2010). Therefore, evaluating changes in DNA methylation in differentiation-related genes during cell differentiation can clarify the developmental toxicity of insecticides. Methyl-CpG-binding domain (MBD) protein 1 is highly associated with DNA methylation (Hendrich and Tweedie 2003), and the MBD domain binds to methylated CpGs (Baubec et al. 2013, Du et al. 2015), thus representing a powerful target for evaluating DNA-methylating effects.

In this study, we developed enhanced green fluorescent protein (EGFP)-MBD1-nls mouse ESCs with a high-content platform. We evaluated the DNA-methylating effects of insecticides to understand the interactions between DNA methylation and cell differentiation. Both global and gene-specific DNA methylation of cell differentiation-related genes during the cell differentiation was evaluated after exposure to various insecticides such as permethrin, carbaryl, DDT, and imidacloprid. This research provided novel insights into the role of DNA methylation in the developmental toxicity of insecticides.

## Material and Methods

### Chemicals

Dimethyl sulfoxide (DMSO) was obtained from Sigma-Aldrich (St. Louis, MO, USA), and 5-aza-2′-deoxycytidine (5-aza-dc), acetamiprid (ACE), imidacloprid (IMI), permethrin, carbaryl, *p,p*′-DDT, and *o,p*′-DDT were obtained from Wako Pure Chemical Industries, Ltd. (Osaka, Japan). DMSO was used as the primary solvent, with solutions further diluted in cell culture medium prior to use. The final concentration of DMSO in the medium did not exceed 0.1% (v/v).

### Cell culture

Mouse ESCs with EGFP-MBD-nls incorporated in their nuclei were provided by the RIKEN Cell Bank (Tsukuba, Japan). The methylated MBD coding sequence confers methylated DNA-specific binding. EGFP is fused to the MBD1 binding domain linked to MBD-nls to yield EGFP-MBD-nls(Kobayakawa et al. 2007). EGFP-MBD-nls cells were cultured on 0.1% gelatin-coated 60-mm dishes or 48-well plates (1 × 10^4^ cells/well). ESCs were cultured in leukemia inhibitory factor (LIF)-supplemented Dulbecco’s modified Eagle’s medium (DMEM) containing 10% fetal bovine serum (Hyclone, Logan, UT, USA) at 37°C under 5% CO_2_ in humidified air. Cells were induced to differentiate via exposure to LIF-supplmented or withdrawed DMEM containing 5-aza-dc, permethrin, carbaryl, ACE, IMI, *p,p*′-DDT, or *o,p*′-DDT (1 × 10^−8^ to 1 × 10^−6^ M) in DMSO for 48 h (Figure 1A).

**Figure 1.**
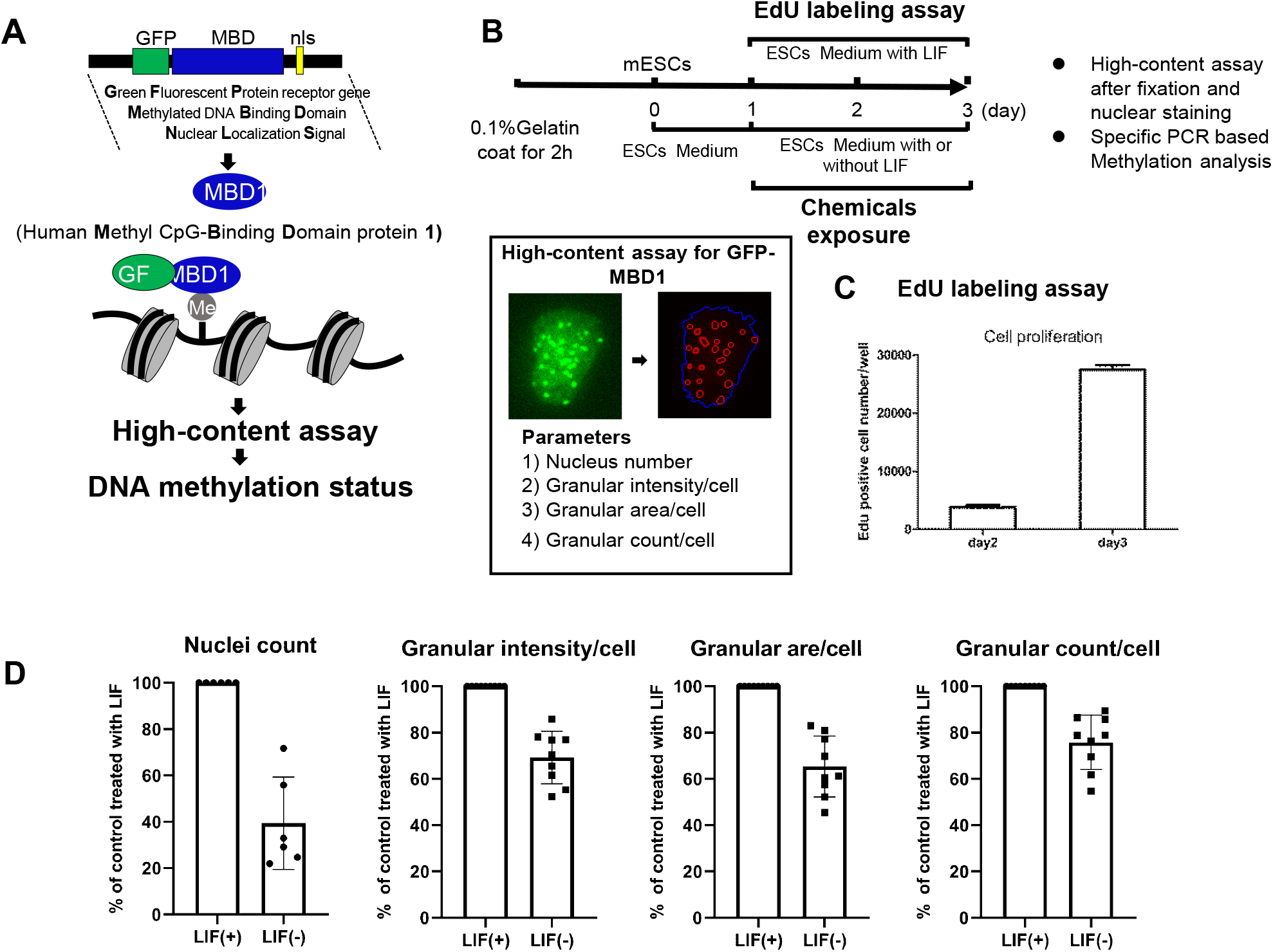
Schematic overview of the detection assay for DNA methylation in the experimental procedure. (A) A depict vector shows that the methyl CpG-binding protein (MBD) and nls coding sequence for human MBD1 confers methylated DNA-specific binding. (B) The high-content assay for GFP-MBD1 determines three phenotypic aspects of the heterochromatin structure, namely the granule staining intensity, area, and count in the nuclei of mouse embryo stem cells (ESCs). (C) EdU cell proliferation assay in undifferentiated status. (D) Effects of LIF withdraw on ESCs in the high-content assay for GFP-MBD1.

### Immunofluorescence

Immunostaining with 4,6-diamidino-2-phenylindole (DAPI) (Life Technologies, Carlsbad, CA, USA) was performed after 48 h of exposure to reagents (Figure 1A). The differentiated cells were fixed with 4% paraformaldehyde for 15 min and then treated with 0.1% Triton X-100 (Sigma-Aldrich) for 30 min. After incubation with 1% bovine serum albumin in phosphate-buffered saline for 30 min at room temperature, cells were incubated with DAPI for 15 min at room temperature.

### Image acquisition and analysis

Microphotographs were obtained using an Olympus LV1200 high-performance laser-scanning microscope (Olympus Optical, Tokyo, Japan). To quantify variations in cellular and subcellular components, immunofluorescent images (four fields per each well of a 48-well plate) were obtained using an automated IN Cell Analyzer 1000 (GE Healthcare UK Ltd., Buckinghamshire, UK) and a ×10 objective. Fluorescence emissions were recorded separately for blue (535-nm) channels. In addition, nuclei (*n* = 100) were counted in each well for confocal image analysis.

The localization of hypermethylated centromeric heterochromatin reflects the nuclear organization and global patterns of DNA methylation (Kobayakawa et al. 2007). To localize centromeric heterochromatin and heterochromatin granule numbers, the granule intensity and area were quantified using a high-throughput assay. Mouse ESCs were cultured in glass-bottomed dishes in parallel to observe heterochromatin variations directly (Figure 1B).

Cells were analyzed using image analysis algorithms generated using the IN Cell Developer Tool Box software (v1.7) in four steps: “nuclear segmentation,” “cell segmentation,” “granular segmentation,” and “measure nodes.” Because a vast array of parameters can be calculated, we selected the following parameters to characterize the DNA methylation status: intensity, area, and number of nuclear heterochromatin granules. Data are expressed as the mean per cell divided by the value for the control. The control values were set at 100%.

### EdU assay

EdU labeling assay was performed with Click-IT EdU Alexa Fluor 555 Imaging Kit (Thermo Fisher Scientific, Japan). Briefly, cultured cells were stained with 10 μM EdU for 24 h at 37°C. After staining, the cells were fixed with 4 % paraformaldehyde for 15 min and permeabilized with 0.5 %Triton X-100 for 20 min at room temperature. After that, the cells were incubated with Click-IT reaction cocktail for 30 min according to according to manufacturer’s instructions followed by Hoechst 33342 staining.

### Methylation PCR analysis

DNA methylation was evaluated using an EpiTect Methyl II DNA Kit provided by Qiagen (Chatsworth, CA, USA) according to the manufacturer’s instructions. In brief, 1 μg of genomic DNA from IMI-treated (1 × 10^−7^ M) mouse ESCs was treated with or without methylation-sensitive enzyme, and then the remaining DNA was mixed with SYBR Green qPCR Master mix for qPCR. The EpiTect Methyl II Signature PCR Array profiles the methylation status of a panel of 22 genes: cell cycle, Brca1, Ccnd2, Cdkn1c, Rassf1, and Cdkn2a; cell apoptosis, Dapk1, Trp73, Fhit, and Apc; cell adhesion, Cdh1 and Cdh13; cell differentiation, Gstp1, Igf2, Vhl, Wif1, and Sfrp1; DNA damage, Mlh1; and tissue development, Timp3, Rarb; Others, Socs1, Pten, and Prdm2. Real-time PCR was performed using an ABI PRISM 7000 Sequence Detection System (Applied Biosystems, Foster City, CA, USA), and the results were analyzed with Δ Δ T and calculated as the methylated percentage in CpG island of individual genes.

### Statistical analysis

Quantitative data are expressed as the mean percentage of the control value ± standard deviation of at least three independent experiments. Analyses were performed using IBM SPSS statistics (IBM Corp., Armonk, NY, USA). Statistical significance was determined using one-way ANOVA for pairwise comparisons. Differences were considered statistically significant when *P* < 0.01.

## Results

### Effect of LIF treatment on EGFP-MBD-nls cells

EdU proliferation assay were performed to investigate the growth status after mouse ESCs post seeding, significantly EdU-positive cells were observed on day 3 (Figure 1C) indicating the cell proliferation of undifferentiated cells. We stimulated the differentiation of mouse ESCs through LIF withdrawal for 72h before DNA methylation assays (Figure 1B). The nuclei number indicating cell viability decreased after withdrawing LIF, and granular intensity reduced may indicate the hypomethylation during the cell differentiation (Figure 1D).

### Effect of 5-aza-dc on EGFP-MBD-nls cells

The DNA-demethylating agent 5-aza-dc is a common anticancer drug that induces DNA demethylation and subsequent activation of tumor-suppressor gene expression (Takebayashi et al. 2001). We applied 5-aza-dc as a model compound to evaluate DNA methylation using the high-content approach. Images of DAPI-stained nuclei were acquired via fluorescence microscopy. The images illustrated that the EGFP-MBD-nls product localized specifically to nuclei and hypermethylated centromeric heterochromatin, which exhibited green fluorescence as granules (Figures 2A).

**Figure 2.**
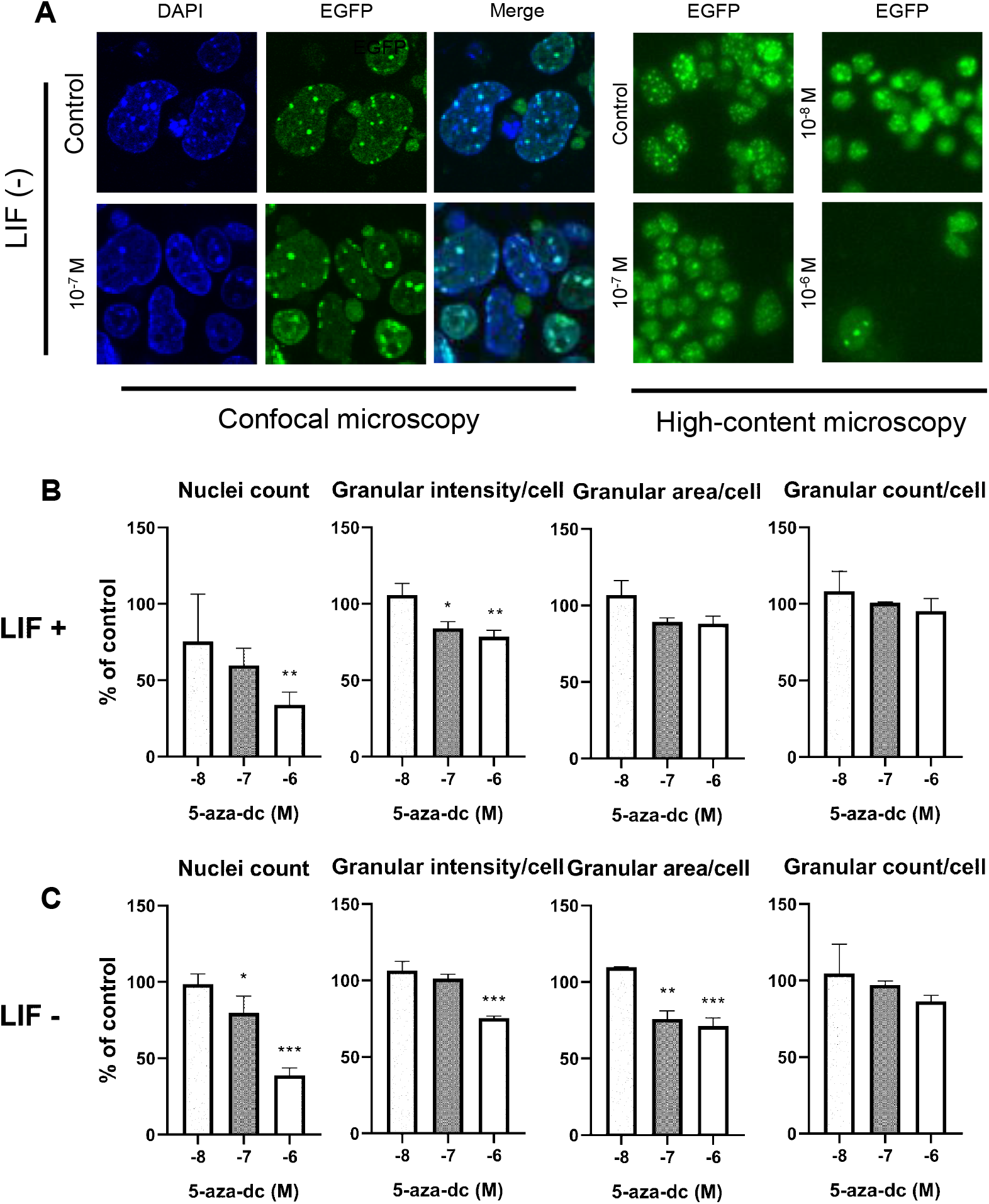
Concentration-dependent effects of 5-aza-2′-deoxycytidine (5-aza-dc) on mouse ESCs. Mouse ESCs were treated with 1 × 10^−8^ to 1 × 10^−6^ M 5-aza-dc for 48 h. (A) Images were taken using an confocal microscopy and a fluorescent microscopy. Graphs showed that the nucleus count, heterochromatin granular intensity per cell, granular area per cell, and granule counts were assessed with LIF (B) or without LIF (C). Data are expressed as the mean of neurite length per cell ± the standard error of three independent experiments. *, P < 0.05; **, P < 0.002; ***, P < 0.001 between the treatment and control groups.

Nuclei count was reduced to 77 (P < 0.05) and 49% (P < 0.05) of the control level by treatment with 1 × 10^−7^ and 1 × 10^−6^ M 5-aza-dc with LIF, respectively. We analyzed granular staining intensity to investigate the effect of 5-aza-dc on methylation, as there is a strong association between the EGFP signal intensity and methylation status in this model system. The granular intensity decreased after exposure to higher concentrations of 5-aza-dc (1 × 10^−6^ M) (P < 0.05), consistent with its demethylation activity (Figure 2B).

Nuclear organization changes are associated with cell differentiation such that changes in the heterochromatic granule count can reflect the differentiation status (Kobayakawa et al. 2007). To evaluate the effect of each chemical on DNA methylation after cellular differentiation, we analyzed granule numbers and areas. After 48 h of treatment, 5-aza-dc (1 × 10^−6^) substantially decreased granular area (*P* < 0.05), and LIF withdrawal decreased the granular area (Figure 2C) may resulting from the cell differentiation. The results suggest that 5-aza-dc simultaneously affects ESC differentiation and induces DNA demethylation.

### Effect of neonicotinoids on EGFP-MBD-nls ESCs

Neonicotinoids have been widely applied as emerging insecticides, but they carry high risks of neurotoxicity in organisms (Solomon and Stephenson 2017). We selected the popular neonicotinoids ACE and IMI for DNA methylation evaluation. ACE exposure reduced nulcei count slightly (Figure 3A). The granular intensity increased to 115 and 128% of control levels after treatment with ACE (1 × 10^−7^ M) (Figure 3A) and IMI (1 × 10^−7^ M) (Figure 3A), respectively, indicating their DNA-methylating effects. After LIF withdrawal, the granular intensity increased to 125 and 145% of control levels after treatment with ACE (1 × 10^−7^ M) (Figure 3B) and IMI (1 × 10^−7^ M) (Figure 3B), respectively, indicating that DNA methylation increases during cell differentiation.

**Figure 3.**
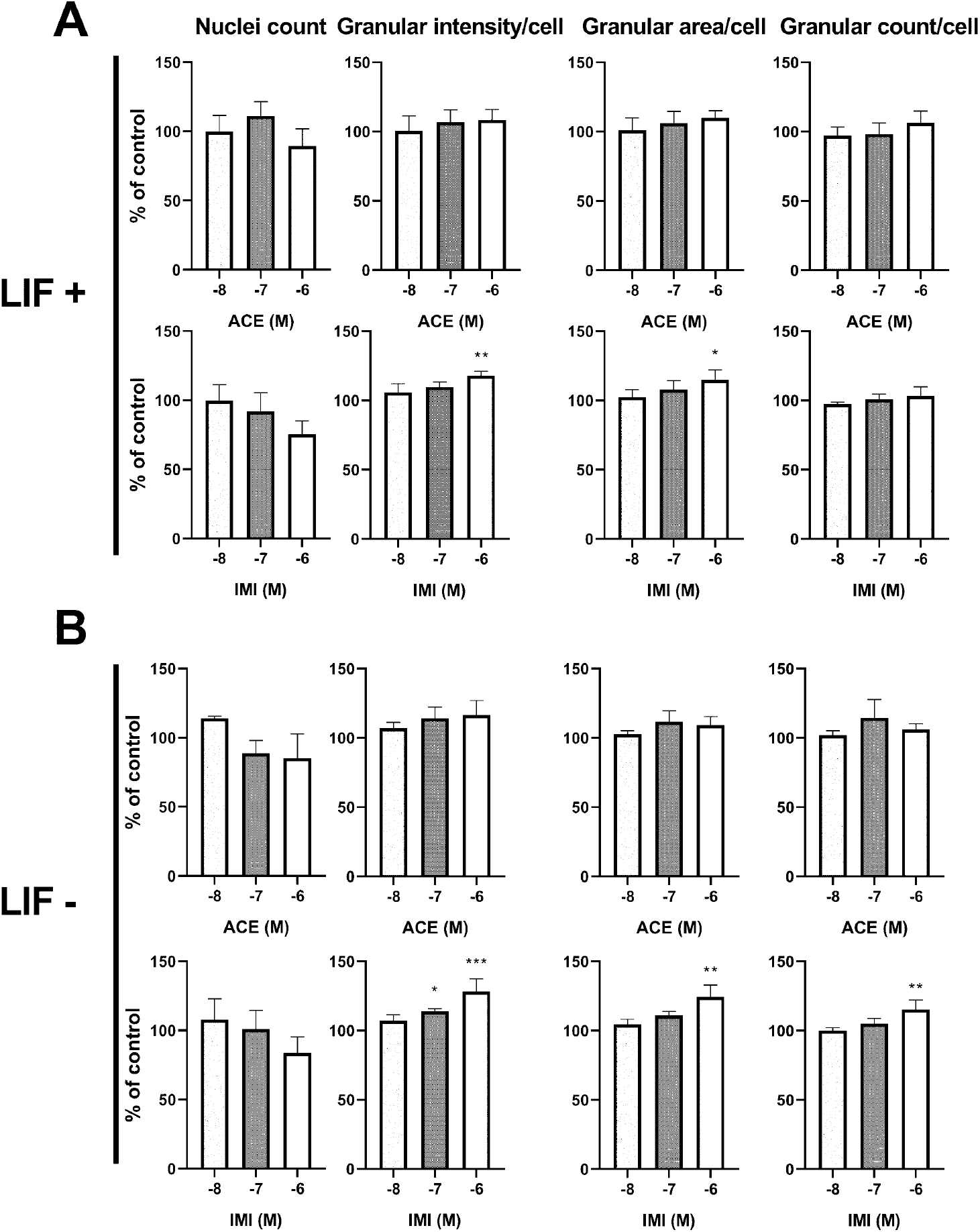
Concentration-dependent effects of acetamiprid (ACE) and imidacloprid (IMI) treatment on mouse ESCs. Mouse ESCs were treated with 1 × 10^−8^ to 1 × 10^−6^ M ACE or IMI for 48 h with LIF (A) or without LIF (B). Nucleus counts, heterochromatin granular intensity per cell, granular area per cell, and granular counts per cell were assessed. Data are expressed as the mean per cell ± the standard error of the mean of three independent experiments. *, P < 0.05; **, P < 0.002; ***, P < 0.001 between the treatment and control groups.

### Effect of IMI on the DNA methylation of specific genes

Alterations in the DNA methylation of functional genes are associated with the development of various cells, and this process can also lead to various diseases. IMI-induced changes in the methylation status of 22 gene promoters were investigated via quantitative real-time PCR in this study. Table 1 shows the DNA methylation status of each gene after exposure to IMI at the early stage of cell differentiation. LIF withdrawal in control cells increased the DNA methylation of most genes by 1.03–12.8-fold. In particular, the DNA methylation of the Ccnd2, Fhit, and Timp3 genes was dramatically increased by 12.80-, 8.38-, and 10.53-fold, respectively. After treatment with IMI, the DNA methylation of all genes decreased dramatically in the undifferentiated stage, but the DNA methylation of the Apc and Gstp1 genes increased in the differentiation stage after IMI exposure. After cell differentiation exposed to IMI, DNA methylation was dramatically increased in the Rassf1, Apc, Gstp1, Timp3, and Pten genes; conversely, the Cdkn2a, Dapk1, Cdh1, Mlh1, and Rarb genes exhibited dramatic decreases in DNA methylation.

**Table 1.** Methylation status of cell proliferation and apoptosis associated genes that were exposed to Imidacloprid (IMI) at 10^−7^ M in the early stage of cell differentiation.

| Biological process in<br>gene ontology | Genes | DNA methylation (%) |  |  |  |
| --- | --- | --- | --- | --- | --- |
|  |  | LIF (+) |  | LIF (-) |  |
|  |  | Control | IMI | Control | IMI |
| Cell cycle | Brca1 | 1.44 | 0.21 | 1.89 | 0.14 |
|  | Ccnd2 | 0.05 | 0.04 | 0.64 | 0.13 |
|  | Cdkn1c | 0.57 | 0.23 | 0.6 | 0.07 |
|  | Rassf1 | 1.45 | 0.22 | 3.52 | 1.75 |
|  | Cdkn2a | 7.82 | 1.82 | 6.7 | 0.77 |
| Cell apoptosis | Dapk1 | 11.31 | 1.74 | 11.4 | 0.41 |
|  | Trp73 | 0.8 | 0.22 | 2.2 | 0.25 |
|  | Fhit | 0.24 | 0.25 | 2.01 | 0.44 |
|  | Apc | 0.42 | 0.19 | 1.6 | 3.16 |
| Cell adhesion | Cdh1 | 15.85 | 2.5 | 25.77 | 1.36 |
|  | Cdh13 | 0.37 | 0.17 | 1.19 | 0.2 |
| Cell differentiation | Gstp1 | 0.42 | 0.08 | 1.03 | 2.54 |
|  | Igf2 | 0.32 | 0.08 | 1.46 | 0.4 |
|  | Vhl | 2.7 | 0.72 | 3.11 | 0.21 |
|  | Wif1 | 0.83 | 0.5 | 2.62 | 0.48 |
|  | Sfrp1 | 1.57 | 0.42 | 1.61 | 0.57 |
| DNA Damage | Mlh1 | 6.81 | 0.83 | 11.61 | 0.35 |
| Tissue development | Timp3 | 2.38 | 0.26 | 25.05 | 1.1 |
|  | Rarb | 9.06 | 3.02 | 18.86 | 0.8 |
| Others | Socs1 | 0.95 | 0.23 | 1.9 | 0.29 |
|  | Pten | 0.2 | 0.01 | 0.77 | 0.61 |
|  | Prdm2 | 0.15 | 0.02 | 0.38 | 0.05 |

The equivalent amount of DNA from 3 wells were pooled and used for methylation PCR analysis in single value calculated as the methylated percentage in CpG island of individual genes.

### Effects of other insecticides on EGFP-MBD-nls ESCs

To investigate the effects of other insecticides on DNA methylation, we assessed DNA methylation following treatment with permethrin, carbaryl, *p,p*′-DDT, and *o,p*′-DDT using the same experimental procedure applied for 5-aza-dc (Figure 4). The granular intensity, area and count increased after treatment with carbaryl and *o,p*′-DDT. *p,p*′-DDT and permethrin decreased such three parameters at high concentrations. These results indicate that *o,p*′-DDT slightly increased DNA methylation, whereas *p,p*’-DDT had no effect. Further *o,p*′-DDT at the highest concentration increased cell proliferation of GFP-MBD-nls cells.

**Figure 4.**
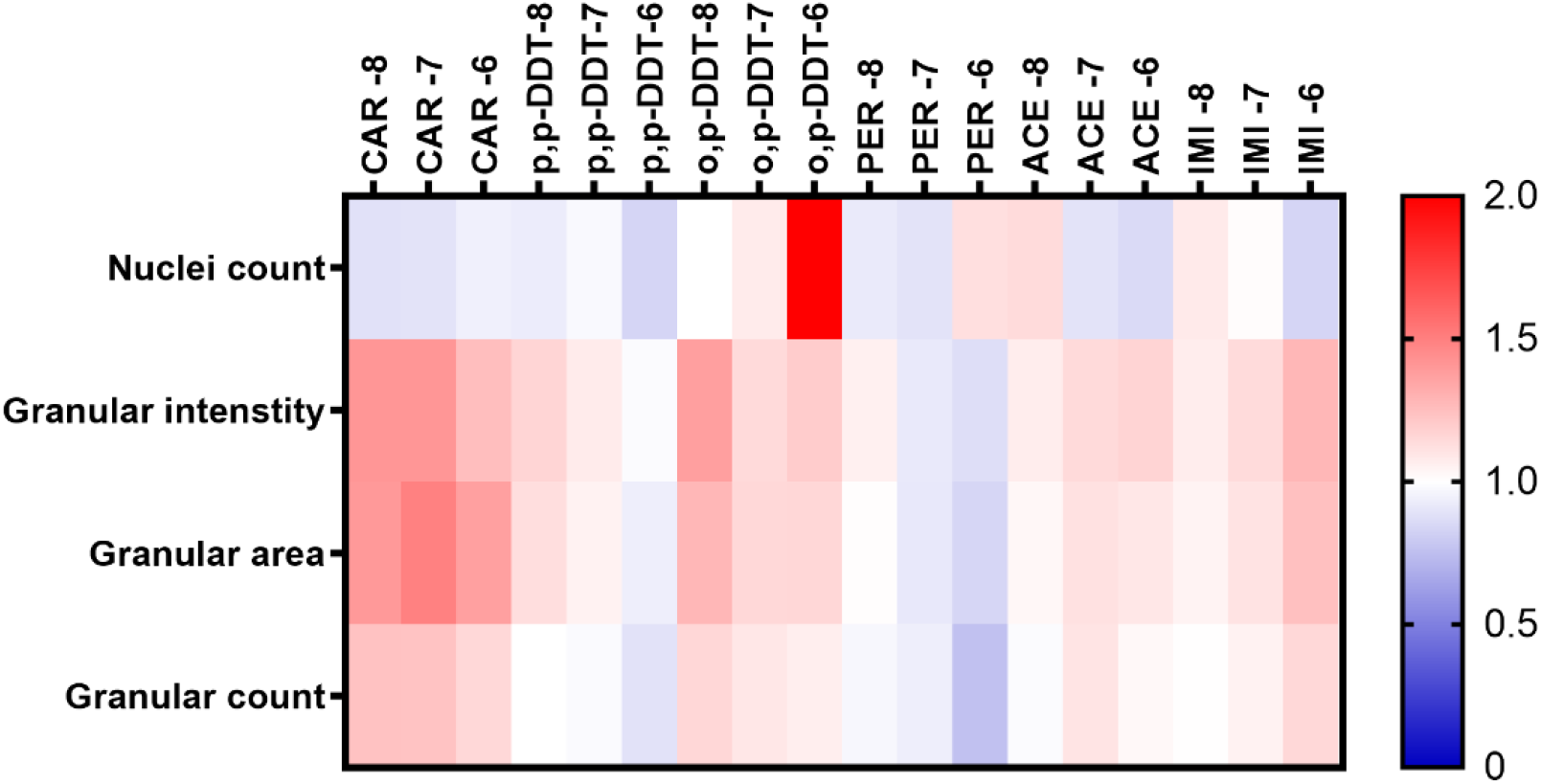
Heatmap of the effects of other insecticides on mouse embryonic stem cells. DNA methylation following treatment with permethrin (PER), carbaryl (CAR), p,p′ -DDT (p/p-DDT), o,p′ -DDT (o/p-DDT), acetamiprid (ACE) or imidacloprid (IMI) at a concentration of 1 × 10^−8^ (–8), 1 × 10^−7^(–7), or 1 × 10^−6^ M (–6) of was examined using the same experimental procedure described or Figure 1. Red and blue in the heatmap indicate increased and decreased DNA methylation, respectively, compared with the control findings.

## Discussion

In the present study, we investigated the effects of several insecticides on global DNA methylation at the early cell differentiation process. We firstly introduced a high-content platform for evaluation of global DNA methylation using mouse ESCs. Our results demonstrated that the insecticides exposure induced global DNA methylation especially in cellular differentiated stages, which demonstrate possible contribution of their DNA methylation to development toxicity.

MBD1 has been well applied as a biomarker to evaluate the global DNA methylation through genetic or proteomic approaches (Badran et al. 2011, Zhang et al. 2012). In EGFP-MBD-nls ESCs, nuclear granules exhibiting MBD1 signals reflect the aggregation of centromeric or pericentric heterochromatin, thus illustrating the DNA methylation status (Kobayakawa et al. 2007). Therefore, the granular intensity, count, and area were assayed as indicators. ESCs exposed to 5-aza-dc exhibited shrinkage or disappearance of these granules, indicating that granule variation was readily characterized by phenotypic profiling. 5-aza-dc prevents DNA methyltransferase 1 from replicating DNA during cell division, thereby decreasing DNA methylation (El Kharroubi et al. 2001). It thus has a dramatic effect on chromosomes, leading to an open chromatin structure (Haaf et al. 2003). EGFP-MBD-nls binds to methylated DNA located in DAPI-stained heterochromatin, which then emits green fluorescence. LIF was used to maintain the pluripotent state and support cell proliferation of mouse ESCs (Cherepkova et al. 2016). After exposure to 5-aza-dc, heterochromatin decreases in both number and intensity because of DNA demethylation (Figure 2A). Multiple HCS parameters correspondingly exhibited decreases in granular intensity, area, and counts (Figure 2). These results are consistent with other approaches using genome-wide analyses of DNA methylation (Takeshima et al. 2015, Wang et al. 2009, Zhang et al. 2004).

Pesticides are widely used and readily detected in the environment, thus posing risks to human health (Weichenthal et al. 2012). Neonicotinoids, which have been increasingly used in recent decades, induce developmental neurotoxicity, reproductive defects, and embryo toxicity through various toxicity pathways based on both in vitro and in vivo results (Babelova et al. 2017, Gu et al. 2013, Kimura-Kuroda et al. 2012). However, the epigenetic effects of neonicotinoids such as DNA methylation on developmental toxicity are not well studied. According to our results, neonicotinoids induced global DNA methylation, and IMI had greater effects than ACE (Figure 3). DNA methylation takes an important role to control of stem cell differentiation (Huang and Fan 2010, Takizawa et al. 2001). Evidence indicated that DNA methylation increases globally during mouse ES cells differentiation (Zhao et al. 2014). Interestingly, exposure with Imidacloprid at high concentration (10^−6^ M) induced a higher hypermethylation (P<0.05) indicating its possible effects on cell differentiation in DNA methylation patterns.

To evaluate developmental toxicity, cellular markers including differentiation-, cell cycle-, and apoptosis-related genes were selected as endpoints (Jung et al. 2015). Therefore, investigation of gene-specific DNA methylation can better clarify the role of DNA methylation in developmental toxicity after insecticides exposure. In this study, 22 genes related to various biological processes including cell cycle progression, apoptosis, adhesion, differentiation, DNA damage response, and tissue development were selected. Based on the results (Table 1), all 22 genes exhibited hypermethylation during cell differentiation after withdraw LIF. Most of the genes are considered inhibit regenerative capacity by controlling cellular processes such as cell cycle progression in stem cells (Pardal et al. 2005). Therefore, their DNA methylation status can provide novel insights for developmental toxicity evaluations in stem cells. Among these genes, the DNA methylation of the Ccnd2, Fhit, and Timp3 genes increased dramatically. Cyclins and cyclin-dependent kinases have the most important roles in checkpoint control in the cell cycle (Tsihlias et al. 1999). The differentiation of ESCs can be regulated through hypermethylation of these genes. However, the DNA methylation of these genes was decreased dramatically after exposure to IMI, indicating developmental toxicity induced by changes in DNA methylation patterns.

DDT, an organochlorine insecticide, has been detected in soil throughout the world (Daly et al. 2007). It was reported that DDT can target constitutive active/androstane receptor and ERα, stimulating cell cycle progression in mouse liver and MCF-7 cells (Kazantseva et al. 2013, Qin et al. 2011, Tsang et al. 2017). In our study, *o,p*′-DDT but not *p,p*′-DDT increased the proliferation of GFP-MBD-nls cells in our study. Recent research demonstrated that a phytoestrogen enhances mESC self-renewal through upregulating core pluripotency transcription factors mediated by ERα, therefore, *o,p*′-DDT may increase cell proliferation via ERα pathways (Tsang et al. 2017). DDT induces DNA methylation through downregulation of the P53 and P16 genes in the rat liver (Kostka et al. 2014). Our results indicate that *o,p*′-DDT slightly increased DNA methylation, whereas *p,p*′-DDT had no effect (Figure 5). IMI and carbaryl induce neurotoxicity by inhibiting acetylcholine esterase (Lee et al. 2015, Millar and Denholm 2007). However, little is known regarding their effects on DNA methylation, although our results indicate that both agents induce DNA methylation with relatively high efficiency. Few studies have examined the effects of pesticides on DNA methylation. Our HCS method may improve our understanding and permit the rapid detection of pesticide-induced DNA methylation.

In conclusion, our present study is the first to evaluate DNA methylation induced by insecticides through detecting MBD using its fluorescence signature, and we investigated its relationship with developmental toxicity at the early stage of cell differentiation through evaluation of gene-specific DNA methylation. Our results indicate that neonicotinoids influence global DNA methylation, and IMI may lead to developmental effects through the DNA methylation of specific genes related to diverse biological processes. Carbaryl and *o,p*′-DDT may induce DNA methylation, whereas permethrin induced slight DNA hypomethylation, potentially through a different mechanism that should be deeply researched. Our present study provided a promising method for evaluating global DNA methylation and provided a basis for examining the relationship of DNA methylation with developmental toxicity.

## Acknowledgements

We thank Dr. Yang Zeng for the technical support for high-content screen.

